# Protein Engineering with Lightweight Graph Denoising Neural Networks

**DOI:** 10.1101/2023.11.05.565665

**Authors:** Bingxin Zhou, Lirong Zheng, Banghao Wu, Yang Tan, Outongyi Lv, Kai Yi, Guisheng Fan, Liang Hong

## Abstract

Protein engineering faces challenges in finding optimal mutants from the massive pool of candidate mutants. In this study, we introduce a deep learning-based data-efficient fitness prediction tool to steer protein engineering. Our methodology establishes a lightweight graph neural network scheme for protein structures, which efficiently analyzes the microenvironment of amino acids in wild-type proteins and reconstructs the distribution of the amino acid sequences that are more likely to pass natural selection. This distribution serves as a general guidance for scoring proteins toward arbitrary properties on any order of mutations. Our proposed solution undergoes extensive wet-lab experimental validation spanning diverse physicochemical properties of various proteins, including fluorescence intensity, antigen-antibody affinity, thermostability, and DNA cleavage activity. More than **40%** of ProtLGN-designed single-site mutants outperform their wild-type counterparts across all studied proteins and targeted properties. More importantly, our model can bypass the negative epistatic effect to combine single mutation sites and form deep mutants with up to 7 mutation sites in a single round, whose physicochemical properties are significantly improved. This observation provides compelling evidence of the structure-based model’s potential to guide deep mutations in protein engineering. Overall, our approach emerges as a versatile tool for protein engineering, benefiting both the computational and bioengineering communities.

## 1 Introduction

The functions of wild-type proteins (WT) often fall short of meeting the industrial or biomedical needs, due to their low activity or insufficient stability. Therefore, it is necessary to optimize their function through favorable mutations, *i.e*., alternation of amino acids (AA) at several sites in a given protein. This operation holds particular significance when designing antibodies [1–3] or enzymes [4, 5]. A protein typically comprises tens to thousands of AAs, each belonging to one of the 20 distinct AA types. To optimize a protein’s functional fitness, conventional practice dictates a greedy search within the local sequence space. The process involves changing AA sites to enhance the protein’s properties, rendering a mutant with a higher gain-of-function [6].

Obtaining mutants with high fitness necessitates the alteration of multiple AA sites within the protein. However, this process incurs significant experimental costs due to the astronomical number of potential mutation combinations. Moreover, the widely-present negative *epistatic effect, e.g*., adding two positive mutation sites together in a protein often worsens its functionality compared to any of the two single-site mutants [7], makes the process even more difficult. Thus, there is a growing demand for *in silico* evaluation of the fitness of protein variants toward positive epistatic effect, especially for those deep mutants with multiple sites being mutated. Deep learning techniques have been developed to expedite the discovery of advantageous mutants. For instance, Lu *et al*. [8] harnessed 3DCNN to identify a novel hydrolase with a beneficial single mutation site that increases the speed of polyethylene terephthalate (PET) degradation by 7-8 times at 50°C. Luo *et al*. [9] introduced ECNet to predict functional fitness for protein engineering with an evolutionary context, thereby guiding the engineering of TEM-1 *β*-lactamase to identify variants with enhanced ampicillin resistance. Nguyen *et al*. [10] predicted ΔΔ**G** with deep learning methods in identifying Cas12b nuclease variants that possess higher thermostability.

In this work, we conceptualized the prediction of mutation effects as a denoising problem and introduce ProtLGN, a structure-based deep learning method designed to facilitate favorable mutants. Mutations in natural settings could be viewed as random alternations of AA types, where natural selection dictates that only those mutants that exhibit optimal fitness to the environment survive. Computationally, changing the AA types of a WT can be seen as introducing corruptions to the AA features of the protein. Consequently, the task becomes restoring the perturbed AA sequences to identify protein variants with the highest fitness. This requires the model to understand the construction rules of proteins and infer the correlation of protein sequence/structure and functionality from the training protein data.

However, the scarcity of labeled protein data and the uniqueness of distinct protein families make it challenging to train supervised learning models from a limited amount of experimental data obtained from the examined mutants. As an alternative, researchers often opt to pre-train models to encode protein representations from their sequences and/or structures. The learned embeddings can be utilized for *de novo* protein design [11–15], variant effects prediction [16–19], higher-level structure prediction [20, 21], *etc*. Analogous to coaching a novice in a new field, a model’s learning efficiency is maximized by initially exposing it to a substantial dataset of WT, allowing it to grasp the broader context before delving into specific proteins and their functionalities. The pretraining process does not involve any supervision with particular learning tasks or target labels, and it is usually termed as *self-supervised learning*. Existing protein representation learning methods rely heavily on language models to infer sequence-level AA constructions, such as autoregressive sequence encoding [12, 20, 22, 23] and multiple sequence alignment (MSA) [24–26]. Both categories require substantial computational resources, such as hundreds of GPU cards, and massive data mining from millions of proteins. The resource-intensive nature hinders model revisions from a few labeled data, and the latter learning requirement conditions effective model training on substantial volumes of high-quality protein data. To tackle this issue, ProtLGN employed roto-translation equivariant graph convolutions to analyze the microenvironment of AAs at the structure level, considering the physicochemical properties of its spatially closed AA neighbors. This lightweight geometric encoding strategy embeds the structural and biochemical properties of proteins efficiently, which eliminates the need for resource-intensive techniques that have been widely applied in protein language models.

We further applied ProtLGN on six different structurally sensitive proteins [7, 27, 28] to engineer specific properties designed for each of them, and validated the results through wet experiments. It includes different-colored fluorescence protein (GFP, EGFP, BFP, and orangeFP), the variable domain of the heavy chain of a nano-antibody targeting growth hormone (VHH antibody), and Argonaute protein from *Kurthia massiliensis* (KmAgo). These proteins underwent modification to achieve specific, practically meaningful goals. We refined the scoring function by public deep mutational scanning (DMS) data on fluorescence intensity on GFP variants [7] and used this scoring function to engineer the fluorescence intensity of GFP, EGFP, BFP, and orangeFP. We also designed single-site mutants on the VHH antibody to enhance its thermostability and binding affinity with antigens with zero-shot predictions. In addition, we combined a batch of positive single mutation sites on KmAgo and discovered strategies for higher-order (2−7) mutants with further enhancement of DNA cleavage activity. The high success rate, with up to 90% of positive rate in property enhancement and eight-fold improvement in catalytic activity, implies the reliability of our proposed ProtLGN in fulfilling the engineering requirements assessed above. Meanwhile, the steadily increased activity with deeper mutations suggests the model’s ability to achieve positive epistatic effects. Moreover, we conducted extensive *in silico* comparisons of ProtLGN with state-of-the-art sequence and structure-based self-supervised deep learning methods on DMS assay prediction and demonstrated that our method achieves the top performance with a significantly reduced model scale.

To validate ProtLGN’s ability to feature protein structures, we additionally conducted an investigation on identifying protein subcellular localization, in which task a protein’s structure plays a pivotal role in determining its functionality.

In summary, ProtLGN merges as a versatile tool for fulfilling a multitude of empirical requirements in protein engineering. It excels in selecting top-ranking single-site mutants for various protein properties without prior knowledge of specific proteins or their functionalities. It could also learn from a small set of labeled data and generalize the discovered patterns to engineer different proteins of similar functionality. More importantly, it is capable of suggesting favorable combinations of single mutation sites that exhibit positive epistatic effects to enhance the performance of higher-order mutants. All the above highlights ProtLGN’s robust predictive performance and broad applicability in protein engineering.

## 2 Results

### 2.1 Model Pipeline

For an input protein, ProtLGN extracts geometry-aware AA-level hidden representations, which can be used to obtain fitness scores for mutants with residue-level differences. The input protein takes the form of a graph, with each node representing an AA. These AA nodes are interconnected with spatially-close neighboring AAs. Node and edge features are established based on local properties (Section 1.1 in SI) and encoded through equivariant graph convolutions (Section 1.2 in SI). For model training, we facilitate a self-supervised prediction task that learns to recover protein sequences from noisy input. This task yields probabilities for each node, indicating their membership being one of the 20 AA types. The delivered probabilities serve as the basis for scoring the fitness of mutations. Therefore, it is imperative to ensure a comprehensive and robust embedding for scoring mutants accurately. To enhance prediction performance, we incorporate biophysics-aware denoising strategies (Section 4.1) and auxiliary learning strategies (Section 4.2.1). Additionally, we establish a few-shot learning pipeline for fine-tuning the model. This pipeline leverages a limited amount of labeled data derived from wet experiments to refine the scoring function tailored to the specific protein template and property of interest (Fig. 1).

**Fig. 1:**
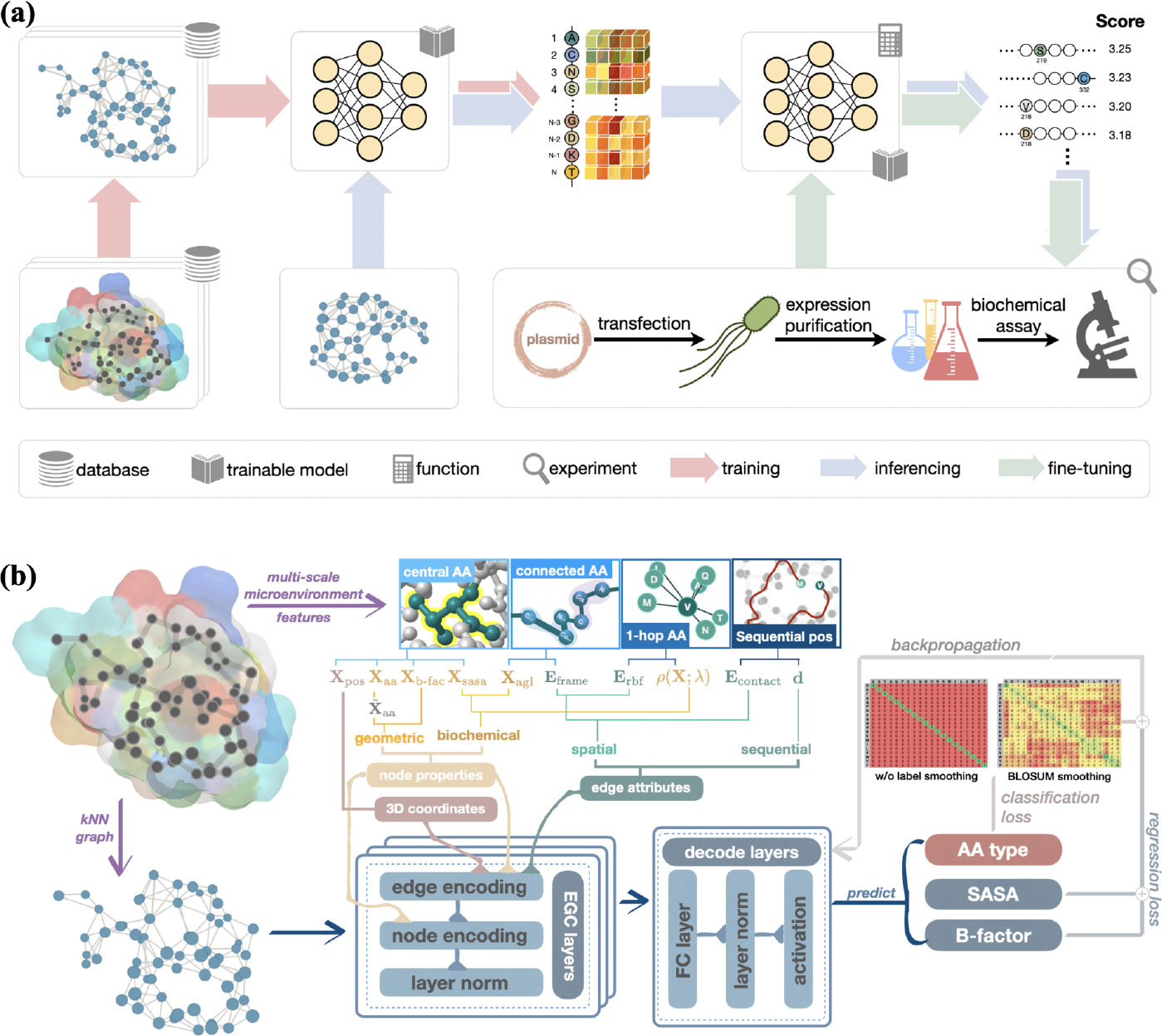
An illustration of the proposed ProtLGN framework for protein engineering. **(a)**. ProtLGN is pre-trained on wild-type proteins for AA-type denoising tasks with equivariant graph neural networks to derive the joint distribution of the recovered AA types (red). For a protein to mutate, the predicted probabilities suggest the fitness score for associated mutations (blue). With additional mutation evaluations from wet biochemical assessments, the pre-trained model can be updated to better fit the specific protein and protein functionality (green). **(b)** Training details for ProtLGN on selfsupervised learning. An input graph is represented by a *k*NN graph with node and edge attributes of the corresponding AAs extracted from multiple scales, as well as nodes’ 3D positions (purple). Wild-type protein graphs with noisy input attributes are fed to EGC layers for rotation and translation equivariant node embedding in the generic protein space (blue).

### 2.2 Properties Enhancement with Biochemical Assessment

We investigated four practical scenarios to comprehensively examine the reliability of ProtLGN for protein engineering. (1) In the first experiment, we assessed the utility of the function-specific fitness scoring function learned from a few labeled GFP variants. The trained scoring function was employed to identify beneficial single mutation sites in the same GFP. (2) In the second experiment, we extended the use of the learned fitness scoring function obtained in (1) to assess its transferability to other fluorescence proteins. Specifically, we aimed to discover ‘the next best’ mutation on EGFP and BFP, both of which share almost the same AAs as GFP with slight variations on 1-3 AAs along the entire sequence [29]. We also examined the same fitness scoring function to orangeFP, coming from a different protein family, characterized by a distinct active region and*∼* 21% sequence identity with GFP. (3) In the third experiment, we applied zero-shot ProtLGN to suggest single-site mutants for the VHH antibody to enhance its binding affinity with antigens and improve its thermostability. (4) The last experiment focused on the search for the optimal combination of positive single mutation sites to enhance properties. We assessed DNA cleavage activity on KmAgo mutants involving changes in 2*−*7 AA sites. Collectively, these wet experiment results across a spectrum of proteins and biological functions affirm that ProtLGN’s fitness predictions on possessing favorable success rates and transferability for single-site mutants and obtaining positive epistatic effects on multi-site mutants.

#### Fluorescence Proteins: Fluorescence Intensity Enhancement with Transferable Fine-Tuned Scoring Function

Fluorescent protein (FP) plays a pivotal role in fields including molecular biology, cellular biology, genetic engineering, and neuroscience [30]. Multicolor FPs enable the concurrent imaging of multiple targets, advancing the study of complex systems [31]. Improving fluorescence intensity in FPs offers advantages for increasing sensitivity for detecting low protein levels, as well as enhancing imaging resolution and signal-to-noise ratio of complex systems.

To customize a scoring function for the fluorescent intensity, we fine-tuned ProtLGN using 1, 000 labeled mutants randomly picked from the deep mutational scanning (DMS) data for GFP [7]. We then ranked the top 10 single-site GFP mutants from all single-site variants that have not been experimentally examined by Sarkisyan *et al*. [7] for wet-lab expression and examination (see more details in Methods). Among these 10 mutants, 5 exhibited higher fluorescence intensity than the WT, with the best-performing mutant achieving fluorescence intensity three times greater than that of the WT (Fig. 2(a)).

**Fig. 2:**
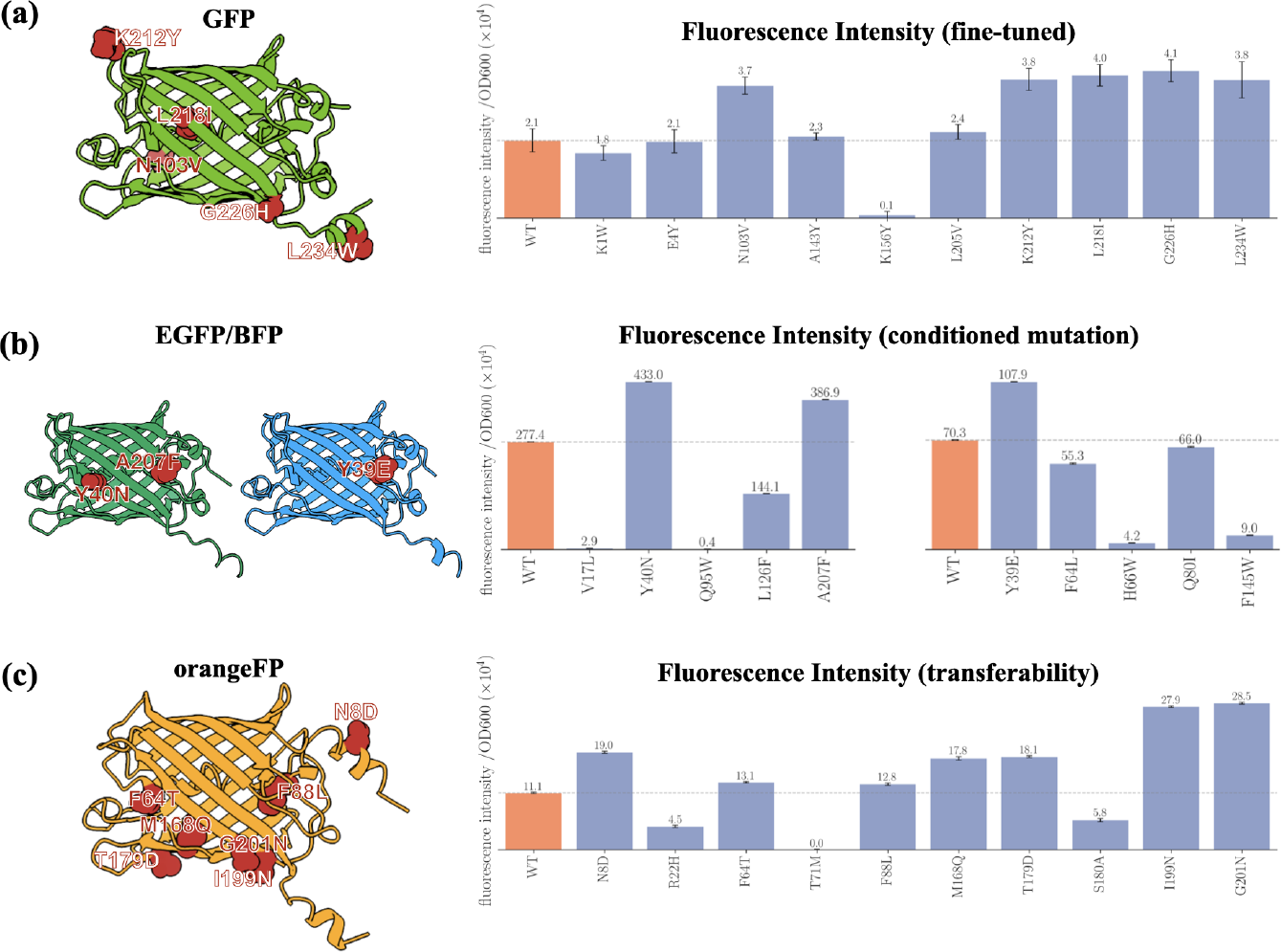
Biochemical results of fluorescence proteins (GFP, EGFP, BFP, and orangeFP) mutants designed by ProtLGN. Left panels: The structure of **(a)** GFP, **(b)** EGFP and BFP, and **(c)** orangeFP. The mutated amino acids with enhanced performance are highlighted in red spheres with annotations. Right panels: The fluorescence intensity by single-site mutants for **(a)** GFP, **(b)** EGFP and BFP, and **(c)** orangeFP. Results from three independent experiments were quantified with the associated standard deviations represented by error bars. Details are presented in Methods and SI.

The learned scoring function is next employed for engineering EGFP and BFP. Both FPs are mutated from GFP with 1*−*3 AAs differences. Particularly, EGFP changed 3 AAs (F64L/S65T/H231L) from GFP, and BFP modified 1 AA (Y145F) from GFP [29]. The same fine-tuned model we trained above was employed again to rank all single-site variants for EGFP and BFP, respectively. For each of the two FPs, the top 5 variants are picked for further wet-lab expression and examination. Two of the modified EGFP exhibited higher fluorescence intensity than the WT, while for BFP, 1 exhibited higher fluorescence intensity than its WT (Fig. 2(b)). This result underscores ProtLGN’s ability to continuously enhance protein functionality when it has been slightly engineered.

Furthermore, we tested the transferability of fine-tuned ProtLGN on a FP from a different family, *i.e*., orangeFP. Both orangeFP and GFP are structurally similar with barrel-like constrictions, but their sequence identity is *∼*21%, and they are fluorescent on different active sites [29]. Same as the previous experiments, we leveraged the fined-tuned ProtLGN to rank the top 10 single-site mutants for orangeFP for wet-lab expression and examination. Impressively, 7 out of the 10 mutants exhibited higher fluorescence intensity than the WT, signifying the model’s robust transferability (Fig. 2(c)).

#### VHH Antibody: Binding Affinity and Thermostability Enhancement by Self-Supervised Learning

The VHH antibody is widely used in medical research and clinical antibody-based drugs. It has been harnessed as an affinity ligand for the selective purification of biopharmaceuticals [32, 33]. Thus, a VHH antibody must undergo mutations to enhance its binding affinity with antigens.

We utilized the pre-trained ProtLGN (learned from*∼*30, 000 unlabeled protein structures, see Section 2.1 in SI) without any experimental data as the input, and selected the top 10 mutants for VHH antibody variants with the highest fitness prediction. We next assessed these 10 mutants for their binding affinity and thermostability through wet experiments. The antibody-antigen binding affinity of the VHH antibody was analyzed by enzyme-linked immunosorbent assay (ELISA) experiments (Fig. 3(a)). Out of the 10 single-site mutants, 4 exhibited a remarkable enhancement in binding affinity, achieving a notable success rate of 40% with the most outstanding mutant achieving an affinity 2.5 times higher than that of the WT. The thermostability of these mutants is measured by differential scanning fluorescence (DSF). As illustrated in Fig. 3(b), 9 mutants exhibited higher melting temperatures (**T**_*m*_) in comparison to the WT, with the highest **T**_*m*_ reaching 45.4 C, 3.4 °C higher than the WT. Intriguingly, 3 mutants displayed a synergistic improvement in both binding affinity and thermostability, which suggests the great potential for ProtLGN-guided mutation design toward extending the lifespan of antibodies while maintaining their binding efficacy.

**Fig. 3:**
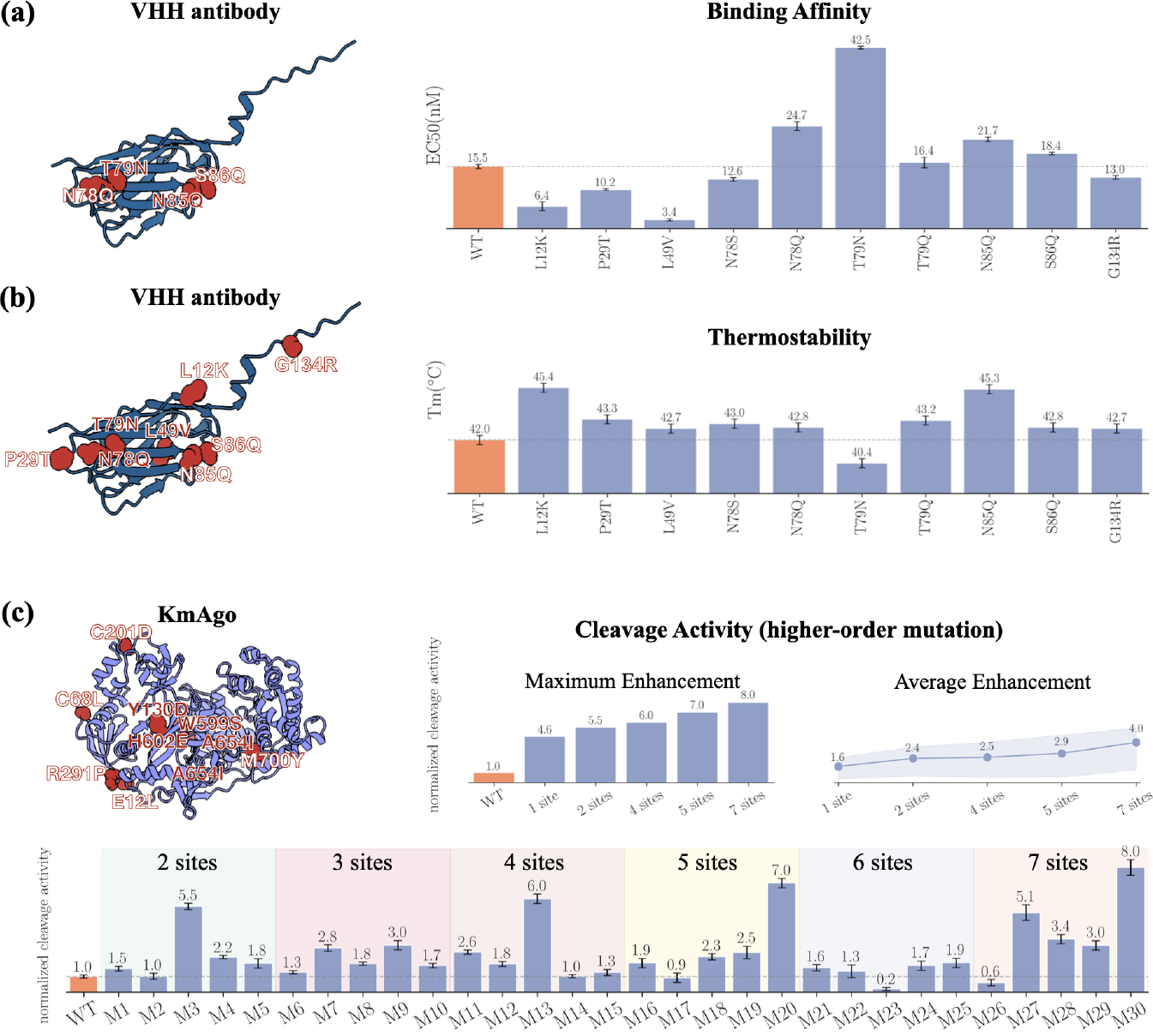
Biochemical results of VHH antibody and KmAgo mutants designed by ProtLGN. **(a)** Left panel: The structure of VHH antibody. Right panel: The binding affinity of VHH antibody and single-site mutants. **(b)** left panel: The structure of VHH antibody. Right panel: The melting temperature of VHH antibody and single-site mutants. **(c)** Upper left panel: The structure of KmAgo. Upper middle panel: The activity of best mutants at different numbers of mutation sites. Upper right panel: The average and standard deviation of the activity of mutants at different numbers of mutation sites. Bottom panel: The cleavage activity of KmAgo and multi-site mutants. The multi-site mutants of KmAgo consist of the combination of sing mutation sites presented in Fig. S19 [39]. The multi-site mutants are presented in Table S9. The mutated amino acids with enhanced performance are highlighted in red spheres with annotations. Results from three independent experiments were quantified with the associated standard deviations represented by error bars. Details of experiments are presented in Methods and SI.

#### Argonaute Protein: Multi-Site Mutants with Single-Site Combinations

Similar to the CRISPR-Cas system, Ago proteins are nucleic acid-guided endonucleases proposed as suitable candidates for various biotechnological applications [34–36]. Among different types of Ago proteins, KmAgo can perform the function near room temperature, which is of great value for developing gene-editing or diagnosis toolkits. However, the application of KmAgo in nucleic acid manipulation encounters challenges due to the low activity of mesophilic Ago proteins [37, 38]. When enhancing protein functionality, singlesite mutants often fall short of achieving the desired function, necessitating the exploration of multi-site mutants for functional evolution. However, due to the negative epistatic effect, achieving deep mutants with enhanced performance often requires enormous experimental efforts and many rounds of iterative search.

Our strategy started from 12 positive single-site mutants with cleavage activities surpassing those of the WT obtained from earlier work [39]. We leveraged ProtLGN to score the combinations of these 12 mutations and select the respective top 5 mutant candidates for 2 to 7 sites. In total, 30 higherorder mutations were picked in a single round to perform wet-lab evaluations. Remarkably, 90% of these mutants exhibited enhanced DNA cleavage activity compared to the WT (Fig. 3(c)). The best mutant is a 7-site mutant, with an 8-fold enhancement on the activity over WT. More importantly, for both the maximum activity enhancement and the average enhancement, higher-order mutants tend to show higher activity than lower-order mutants. This result demonstrates that our model can deliver deep mutants of high gain-of-function and identify the positive epistatic effect when combining single-mutations sites.

### 2.3 Fitness Prediction on Deep Mutational Scanning Benchmarks

To further evaluate the capability of ProtLGN on a broader range of applications, we compared its performance in predicting protein fitness in public DMS datasets with other state-of-the-art self-supervised learning models. Fig. 4(a) compares self-supervised learning baseline methods that analyze proteins with their structure, multiple sequence alignment (MSA), or sequences. We are particularly interested in evaluating property enhancement with higher-order mutants as it has been shown that achieving the desired physicochemical properties frequently requires mutating on multiple sites (Fig. S2). As shown in Fig. 4(a), overall, ProtLGN achieves the top performance with the fewest trainable parameters. This substantial reduction in expense not only facilitates pre-training a general-purpose model through self-supervised learning but also enhances the fine-tuning of protein and property-specific algorithms at later stages when experimental labels become partially available.

**Fig. 4:**
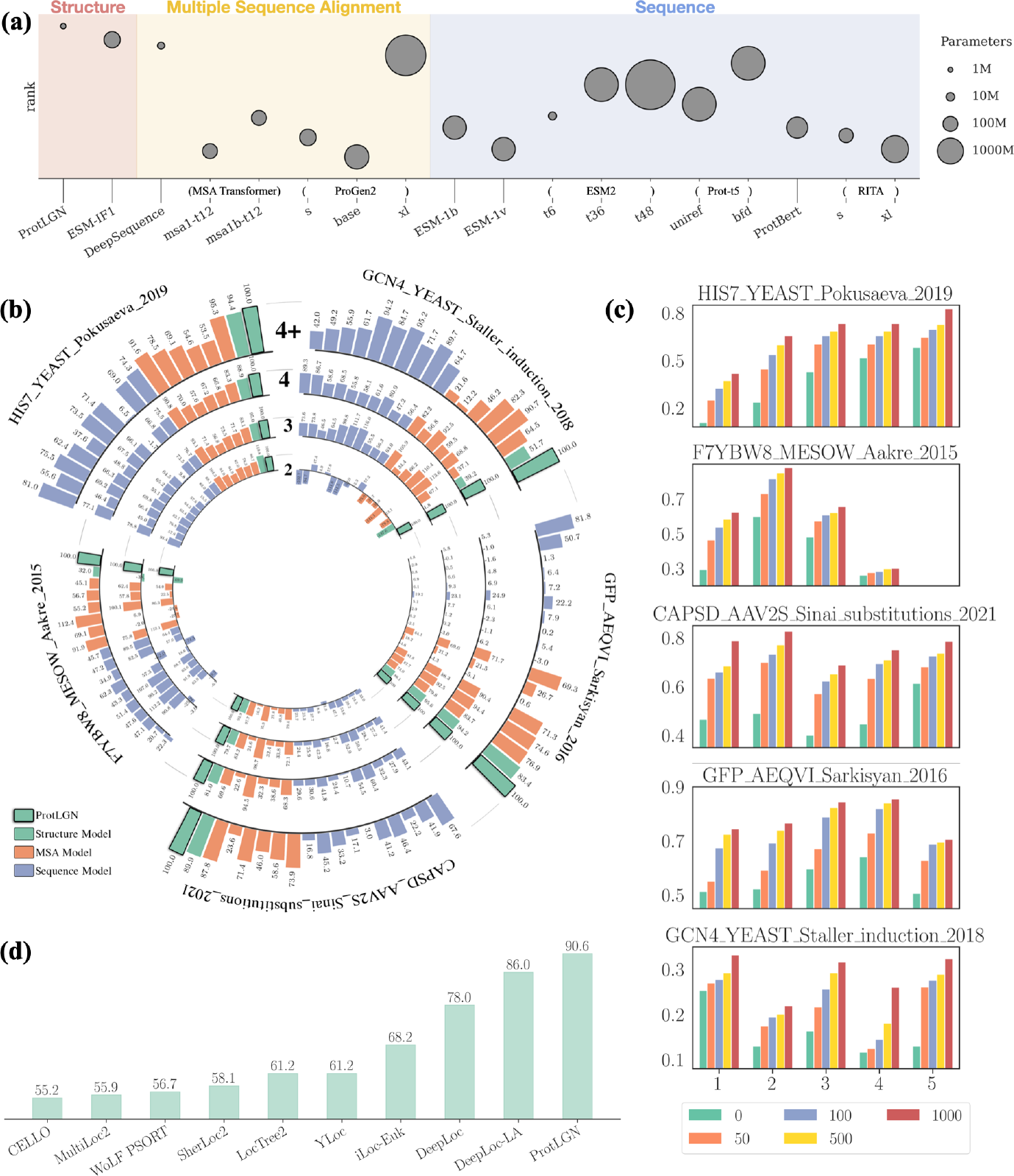
*In silico* evaluation with variant effects prediction and protein subcellular localization prediction. **(a)** Inference efficiency and effectiveness on zero-shot deep learning models. A higher altitude on the *y*-axis indicates the model ranks higher on variant effects prediction. The radius of each ball indicates the number of trainable parameters of the associated model. **(b)** Per-protein performance on the multi-site variant effects prediction with models trained from zero-shot learning tasks. Circles (from inside to outside) represent prediction performance on higher-order mutants changing 2, 3, 4, and *>* 4 AAs. Bars are colored by modeling approaches, including structure-based models (green), MSA-based models (orange), and sequencebased autoregressive models (blue). Values on each bar indicate the percentage of the associated method’s relative Spearman’s correlation with respect to ProtLGN(normalized to 100). **(c)** The performance gain of ProtLGN’s variant effects prediction on higher-order mutations when fine-tuning on additional 100 *−* 1000 randomly picked labeled mutations. **(d)** Classification accuracy on identifying protein subcellular localization on the test proteins.

In Fig. 4(b), we assessed ProtLGN’s fitness prediction performance on five proteins, each with ground-truth scores on at least 4-site mutants (See details in Table. S1). For clearer comparison, we normalize Spearman’s *ρ* against ProtLGN’s performance (standardized to 100%). Overall, we observe a significant outperformance of ProtLGN for higher-order mutations across all five proteins. In contrast, the baseline methods in many cases achieve correlations close to zero or even negative correlations when compared to the ground truth. In the case when a small set of mutant-specific ground-truth scores is available, ProtLGN can be fine-tuned effectively to cater to a specific protein given its lightweight nature. We refined the scoring function using labeled data to better suit the protein and its associated properties (See Section 4.2). For each protein, these 100*−* 1, 000 training samples were randomly picked from the corresponding DMS dataset with an arbitrary number of mutational sites and fitness scores. As displayed in Fig. 4(c) and Fig. S5, a few hundred mutation scores suffice to achieve a notable correlation score.

### 2.4 Protein Subcellular Localization Predictions

Protein subcellular localization (PSL) refers to the specific location of a protein within a cell [40]. In different organelles, proteins possess distinct structural features that align with their individual biological functions, as the functionality of an intracellular protein is intrinsically tied to its structure [41–43]. For instance, different membrane proteins perform distinct functions, such as transport and signaling, and exhibit structural diversity across transmembrane, extracellular, and intracellular domains [44]. Another example is provided by lysozyme, which participates in the immune system through the adoption of alpha/beta protein folds for functionality within the Lysosome/Vacuole [45]. Consequently, achieving accurate PSL prediction hinges on a deep understanding of the functionality and physicochemical characteristics of proteins.

In this context, it is imperative to highlight the significance of employing protein topological information when generating hidden embeddings through a structure-based model like ProtLGN. We followed the construction of [46] for the PSL prediction task and compared the prediction performance of baseline methods reported there. ProtLGN was trained on a set of 9, 366 labeled proteins, with each protein represented by a vector pooled from its AA-level representations. The evaluation was conducted on 2, 738 test proteins to predict whether these proteins exist on 10 possible locations of cells. As displayed in Fig. 4(d), ProtLGN significantly outperforms existing prediction methods in accurately predicting the subcellular localization of proteins. It accomplishes this by comprehensively analyzing the intricate 3D representation of proteins, which encapsulates vital spatial folding information that governs biological functions, such as secondary structure, topological domains, and protein surface characteristics [42]. In contrast, existing baseline methods rely on AA sequences [46] or homology information [47, 48] when predicting PSL. While these methods attempt to identify protein regions that are crucial for subcellular localization, such sequence-based approaches are not as direct as structure-based methods, and thus were outperformed by ProtLGN.

## 3 Discussions and Conclusions

Multi-site mutants are key players in protein engineering, leading the way toward enhanced functionality by aggregating the effects of positive singlesite mutants. Discovering effective multi-site mutants requires the deliberate combination of single-mutation sites exhibiting improved functionality. Traditionally, identifying favorable mutants relied on experimental or experience-based methods, such as rational design, random mutagenesis, and single-site saturation mutagenesis [49, 50]. However, these approaches demand extensive experimental investments to validate functionally enhanced mutants across all potential combinations. This process consumes considerable time and human resources, yet frequently yields a relatively low success rate due to the vast search space. Moreover, multiple iterations of evolution are typically required to identify and integrate beneficial mutations, *e.g*., creating a 10-site mutant often entails at least ten cycles of high-throughput scanning or a greedy search. The presence of mutation interdependencies (epistasis), rooted in the intrinsic physiochemical nature of proteins, gives rise to a discontinuous fitness landscape that masks the mutation routes to global optimization [51].

To counteract the challenge posed by the expansive yet unsmooth search space and the limited number of experimental data, we introduced a lightweight deep learning solution for scoring protein variants’ fitness. This approach leverages the microenvironment of amino acids to efficiently interpret protein construction rules based on a small set of labeled experimental data associated with the biochemical assays of protein variants. The established ProtLGN demonstrates a remarkable ability to transfer knowledge and effectively engineer advantageous mutants with distinct functions across different proteins. Empirically, ProtLGN achieves a favorable success rate of around 40% when engineering single mutation sites in various proteins with distinct functional enhancements, including fluorescence intensity, thermostability, and binding affinity. The exceptional performance of ProtLGN positions it as a promising solution for fulfilling various practical requirements in protein engineering. More importantly, ProtLGN has shown its capability to circumvent the negative epistatic effect with a cost-efficient strategy, which makes it possible to discover advantageous deep mutants through low-throughput screening of single-site mutants. The ProtLGN-guided approach for multi-site mutants on KmAgo achieves superior DNA cleavage activity compared to the performance of low-order mutants by effectively capturing the epistatic effect among mutation sites. Notably, the higher-order mutants are constructed in a single step, drawing from the pool of tens of experimentally characterized single-site mutants.

We attribute the success of the cost-efficient design for single-site and multisite mutants to ProtLGN’s capacity to incorporate the impact of mutations on the protein’s local structure. We also validate this deduction by training ProtLGN on an additional protein classification task, where ProtLGN accurately predicts the localization of the protein within the cell, in which case the structure-relevant factors are highly related to protein functionality. Moreover, ProtLGN is capable of inferring the joint distribution of multiple amino acid sites simultaneously, thus eliminating the need to assume independent mutations – a limitation that characterizes many autoregressive models in finding the conditional score for individual mutations at a single site. Consequently, the performance of ProtLGN in designing amino acid mutations is a pivotal asset in the protein engineering strategy.

## 4 Methods

### 4.1 AA Representation Learning

To encourage the network to capture essential representations of AAs, we integrate various AA characteristics into our pre-training process. We simulate random mutations that occur in nature and introduce noise to node features throughout the training phase. To guide AAs toward higher probabilities of substituting with groups that they commonly transition to in nature, such as polar, charged, and hydrophobic groups, we implement substitution matrices for both noise generation and label smoothing stages.

#### 4.1.1 AA Type Purterbation

The AA type of a node ***x***_aa_ undergoes perturbation, resulting in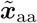with a multinomial noise, *i.e*.,

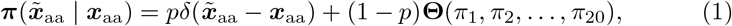

where the confidence level *p* is a tunable parameter that controls the fraction of AAs considered ‘noise-free’. This parameter can also be determined based on prior knowledge regarding the quality of wild-type proteins, reflecting the expected frequency of mutations in nature. For instance, a value of *p* = 1 indicates maximal confidence in the quality of wild-type proteins, resulting in no perturbations to the input AA.

The probability of an AA adopting a specific type is contingent on the defined distribution **Θ**. A straightforward choice for **Θ**, representing the expected frequency distribution of AA types, involves setting them to equal values: *π*_1_ = *π*_2_ = … = *π*_20_ = 0.05. It can also be informed by prior knowledge derived from molecular biology, such as the observed probability density of AA types in wild-type proteins. Another option for defining pair-wise exchange probabilities is through substitution matrices. The impact of different *p* values is illustrated in Fig. S6. A sharp decline at *p* = 1 confirms the efficacy of introducing perturbations to AA types in enhancing fitness prediction performance for both single-site and multiple-site mutations. Additionally, an AA-specific perturbation probability surpasses a random corruption rate in terms of overall performance enhancement. Generally, assigning a moderate value to *p* between 0.4 and 0.7 is suitable for various combinations of learning modules and noise distributions. Based on overall performance, we have set the default value of *p* to 0.6 as the confidence level for our model.

#### 4.1.2 Label Smoothing

Protein sequence alignments play a pivotal role in understanding potential substitutions of one AA for another. To capture these insights, we utilize the BLOSUM62 substitution matrix [52], a 20 *×* 20 square matrix denoted by ***B***, that defines pairwise similarities between AA types based on relative substitution frequencies and chemical characteristics. We implement BLOSUM62 for label smoothing, where AAs reduce penalties for AAs when they mutate to types with high similarity scores in the matrix. The degree of dispersion towards other AA types, as indicated by the off-diagonal regions of BLOSUM62, is controlled by temperature *t*. The transformation from ***B*** to ***B***^*′*^ is achieved through ***B***^*′*^ = *σ*(***B***)^*t*^, where *σ*(·) represents a non-linear operator, such as normalization. Intuitively, increasing the value of *t* tends to align ***B*** more closely with a diagonal matrix, irrespective of the higher confidence level *p* associated with wild-type noise (see Fig. S7). The modified ***B***^*′*^ with different *t*s for defining the label smoothing and perturbation probability is displayed in Fig. S7. In comparison to baseline results involving random AA noise and no label smoothing, applying the BLOSUM62 matrix to label smoothing leads to improved predictions, where a higher temperature yields better overall performance. We also assessed the impact of varying *t*s on AA type perturbation, where a moderate *t* that produces more nuanced noise to AA types is preferred.

### 4.2 Tunable Variant Effects Prediction

#### 4.2.1 Multitask Learning Strategy

During the recovery of the AA site distribution, ProtLGN corrects perturbed AA types and predicts the joint distribution of all AA types, denoted as ***y***_*aa ∈*_ *ℝ*^20^, for mutation scoring. To enhance the expressiveness of the hidden node representation, we employ a multitask learning strategy within the selfsupervised learning module. Two auxiliary tasks are introduced, involving the prediction of ***y***_sasa_ and ***y***_B-fac_. The former task pertains to solvent-accessible surface area (SASA), which significantly influences AA type preferences, while the latter involves B-factors, which are related to the conformations and mobility of neighboring AAs. Both properties are closely connected to AAs and are essential for characterizing individual AAs. Predicting these two features aids in implicitly encoding these characteristics, which substantially enhances prediction performance, irrespective of the selected noise distribution (Fig. S6). While all three learning tasks collectively improve the embedding effectiveness, it is crucial to strike a balance among them and regulate their contributions to the overall loss function:

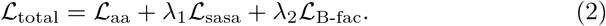

Fig. S8 explores a range of choices for *λ*s and their impact on model training. Since both ***y***_sasa_ and ***y***_B-fac_ share a consistent value scale, we set *λ*_1_ = *λ*_2_*∈ {*0.05, 0.1, 0.2, 0.5, 0.8, 5, 10*}*. All experiments are conducted on zero-shot multi-site mutations with wild-type AA noise (*p* = 0.6). The overall declining trend indicates that accurately predicting ***y***_aa_ remains the model’s primary objective. Notably, a small peak in the scores near *λ* = 0.1 suggests that smaller *λ* values are generally preferred.

Fig. S9 examines the consistency between predicted results and the ground truth for ***y***_*aa*_, ***y***_sasa_, and ***y***_B-fac_. The near-diagonal confusion matrix for predicted AA types compared to ground-truth AA types evaluates ProtLGN’s ability to recover from noisy sequences to the original sequence. For the two regression tasks involving SASA and B-factor predictions, linear regression is applied to fit the true values against the predicted values, yielding estimated coefficients of 0.989 and 1.008, respectively, both with *p*-values *<* 0.001. Besides, Pearson’s correlation coefficients between the predicted and true values of SASA and B-factor are reasonably high at 0.884 and 0.791.

#### 4.2.2 Scoring Function for Fitness Prediction

The recovered AA distribution for the protein to some extent reflects the rational appearance of a naturally selected protein. When investigating an arbitrary protein with a ‘cold-start’ *i.e*., no experimentally tested data initially, we follow [8, 18] and define the *log-odds-ratio* as a substitution for the fitness score, which is directly obtained from probabilities. For *T* -site mutants (*T≥* 1), the fitness score reads

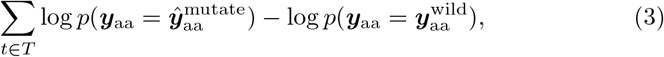

where 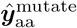 and 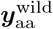 represent the predicted and wild-type AA types, respectively.

Since the recovered AA distribution does not always provide the optimal evolutionary direction for any desired property in protein engineering, it is advisable to fine-tune the designed general-purpose model to better align with protein-specific or task-specific contexts when possible. If the mutation strategies for the protein are partially evaluated, *i.e*., a certain amount of labeled experimental results for mutational assays is available, the predicted probabilities can be transformed into the mutational score of interest using additional fully-connected layers 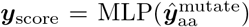 to craft proteinand propertyspecific scoring functions. The training objective is to minimize the disparity between the predicted and actual score distributions, and for this purpose, we employ KL divergence as a measure of dissimilarity.

### 4.3 Protein Preparation

#### VHH Antibody

The gene of the VHH antibody and mutants was cloned into the pET29a plasmid with an N terminal His-tag (Fig. S15). The expression plasmid was transformed into *E. coli* BL21(DE3) cells. A single colony of each recombinant E. coli strain was inoculated into 30 mL LB medium with 50 *μ*g/mL kanamycin for seed culture at 37 °C for 12-16 h. The seed culture was transferred 10 mL to 1 L LB medium with 50 *μ*g/ml kanamycin at 37 °C 220 rpm until the OD600 value got 0.6-0.8. The culture was cooled to 16 °C and then induced with 0.5 mM IPTG for 20-24 h at 16 °C. Cells were harvested from the fermentation culture by centrifugation for 30 min at 4,000 rpm, and the cell pellets were collected for later purification. The cell pellets were resuspended in buffer A (20 mM Na_2_HPO_4_ and NaH_2_PO_4_, 0.5 M NaCl, pH 8.0) and then lysed via ultra sonification. The lysates were centrifuged for 30 min at 12,000 rpm at 4 °C, after which the supernatants were subjected to Ni-NTA affinity purification with elution buffer (20 mM Na_2_HPO_4_ and NaH_2_PO_4_, 0.5 M NaCl, 250 mM imidazole, pH 8.0). The purity of the fractions obtained was analyzed using SDS-PAGE. The fractions containing the purified target protein were combined and desalted using an ultrafiltration unit. The purified protein was then concentrated and stored in buffer A with 10% glycerol at a temperature of -80 °C. The SDS-PAGE of the VHH antibody and its mutants is shown in Fig. S17.

#### Fluorescence Protein

The gene of GFP, eGFP, BFP, and orangeFP and mutants was cloned into the pET28a plasmid (Fig. S14). The expression plasmid was transformed into *E. coli* BL21(DE3) cells. A single colony of each recombinant *E. coli* strain was inoculated into 30 mL LB medium with 50 *μ*g/mL kanamycin for seed culture at 37 °C for 12-16 h. The seed culture was transferred 10 mL to 1 L LB medium with 50 *μ*g/ml kanamycin at 37 °C 220 rpm until the OD600 value got 0.6-0.8. The culture was cooled to 16 °C and then induced with 0.5 mM IPTG for 20-24 h at 16 °C. Cells were harvested by centrifugation for 30 min at 4,000 rpm, and the cell pellets were collected for washing three times. The cell pellets were resuspended in buffer A (25 mM Tris-HCl, 500 mM NaCl, pH 7.4) for testing.

#### KmAgo

A codon-optimized version of KmAgo and mutant gene was synthesized by Sangon Biotech (Shanghai, China). It was cloned into the pET15b plasmid (Fig. S16) to construct pEX-Ago with an N terminal His-tag. The expression plasmid was transformed into *E. coli* BL21(DE3) cells. A 30 mL seed culture was grown at 37 °C in LB medium with 50 *μ*g/mL kanamycin and was subsequently transferred to 500 mL of LB in a shaker flask containing 50 *μ*g/ml kanamycin. The cultures were incubated at 37 °C until the OD600 reached 0.6-0.8, and protein expression was then induced by the addition of isopropylD-thiogalactopyranoside (IPTG) to a final concentration of 0.5 mM, followed by incubation for 16-20 h at 18 °C. Cells were harvested by centrifugation for 30 min at 4,000 rpm, and the cell pellets were collected for later purification.

The cell pellets were resuspended in lysis buffer (25 mM Tris-HCl, 500 mM NaCl, 10 mM imidazole, pH 7.4) and then disrupted using a high-pressure homogenizer at 700-800 bar for 5 minutes (Gefran, Italy). The lysates were centrifuged for 30 min at 12,000 rpm at 4 °C, after which the supernatants were subjected to Ni-NTA affinity purification with elution buffer (25 mM Tris-HCl, 500 mM NaCl, 250 mM imidazole, pH 7.4). Further gel filtration purification using a Superdex 200 (GE Tech, USA) was carried out with an elution buffer. The fractions were analyzed by SDS-PAGE, and fractions containing the protein were flash frozen at -80 °C in buffer (15 mM Tris–HCl pH 7.4, 200 mM NaCl, 10% glycerol). The SDS-PAGE of KmAgo and its mutants are shown in Fig. S18.

### 4.4 Property Assay Evaluation

#### Binding Affinity of VHH Antibody at pH 13.7

The VHH antibody was incubated with 0.5 M NaOH (pH = 13.7) for 24 hours. 96-well plates were coated with growth hormone protein at a density of 5 ng/well at 4 °C overnight. The plates were washed with 1 × PBST three times. Following blocking with 1% BSA in 1 × PBST at 25 °C for 2 h. After washing three times with 1 × PBST, the plate was incubated with serial dilutions of VHH 100 *μ*L/well (1:2, 1:4, 1:8, 1:16, 1:32, 1:64, 1:128, 1:256, 1:512, 1:1024, 1:2048) for 1 h at 25 °C. After washing three times with 0.5% PBST, 100 *μ*L/well HRP (1:5000) were added and incubated at 25 °C for 1 h. The plate was washed with 1 × PBST four times, and a total of 100 *μ*l/well TMB was added and incubated at 25 °C for 15 min in the dark. Finally, 100 *μ*L/well 2 M H_2_SO_4_ was added to stop the reaction, and absorbance was measured at 450 nm (TECAN, Swiss). The log(agonist) vs. response – variable slope (four parameters) curves were analyzed to calculate EC50 which determines the binding affinity of VHH after alkaline treatment. The original data is shown in Fig. S10.

#### Thermostability of VHH Antibody

Each sample containing 2 *μ*M of protein in 1 × PBST was prepared in triplicate and added to PCR tubes. SYPRO Orange dye available as 5000× stock (Sigma-Aldrich) was added just before the measurement of the proteins in an appropriate amount to achieve a final concentration of the dye of 5×. The thermal denaturation of the proteins was monitored by exciting the SYPRO Orange dye at 470 nm and monitoring its fluorescence emission at 570 nm using Q-PCR (Analytikjena, Germany). The baseline correction is used by the Opticon Monitor software available on the PCR instrument. The original data is shown in Fig. S11.

#### Fluorescence Intensity of Fluorescence Proteins

The cells were washed three times in 1 × PBS, and the OD600 of the cultures was determined. Fluorescence was measured at an excitation wavelength of 365 nm, 381, and, 548 nm using an Agilent Synergy H1MF (Biotek) fluorometer to detect emission at 509 nm, 445 nm, and 562 nm for GFP and EGFP, BFP, and orangeFP, respectively. The fluorescence values of three independently grown cultures were averaged and normalized to the density of cells (five gradient densities of cells were used to normalize the fluorescence intensity (OD600 = 0.4-0.6)).

#### Single-Strand DNA Cleavage Activity of Argonaute Protein

For standard activity assays of KmAgo and its mutants, cleavage experiments were performed in a 2:1:1 molar ratio (protein:guide:target). First, 5 *μ*M protein was mixed with a synthetic 1 *μ*M gDNA guide in the reaction buffer (15 mM Tris-HCl (pH 7.4), 200 mM NaCl, and 5 mM MnCl_2_). The solution was then pre-incubated at 37 °C for 20 min. After pre-incubation, 1 *μ*M tDNA, which was labeled with the fluorescent group 6-FAM at the 5 -end and the quencher BHQ1 at the 3 -end, was added to the mixture. The cleavage experiments were performed at 37 °C. All experiments were performed in triplicate, and the fluorescence signals were traced by the quantitative realtime PCR QuantStudio 5 (Thermo Fisher Scientific, USA) with *λ*_ex_ = 495 nm and *λ*_em_ = 520 nm. The results were analyzed by QuantStudioTM Design & Analysis Software v1.5.1. The activities are determined as the slope of the time dependence of fluorescence intensity resulting from the cleavage function before it reaches the plateau [37, 53]. The guide DNA and target DNA used for cleavage are listed in Table S8. Three independent experiments are conducted to determine the cleavage activity. The original data is shown in Fig. S12-S13.

## Supplementary Information

Figures S1-S19, Tables S1-S9.

## Author Contributions

B.Z. and L.Z. conceived and designed the study. B.Z. designed ProtLGN. L.Z. and B.W. performed biochemical experiments. B.Z., O.L., Y.T., and K.Y. performed ProtLGN for different tasks. B.Z. and L.Z. wrote the manuscript. L.H., B.Z., and L.Z. revised the manuscript.

## Acknowledgments

The authors acknowledge the Center for HighPerformance Computing at Shanghai Jiao Tong University for computing resources.

## Additional Information

Correspondence and requests for materials should be addressed to.

## Data Availability

All the data are presented in SI.

## Code Availability

The model implementation can be found at https://github.com/bzho3923/ProtLGN.

